# NBR: Network-based R-statistics for (unbalanced) longitudinal samples

**DOI:** 10.1101/2020.11.07.373019

**Authors:** Zeus Gracia-Tabuenca, Sarael Alcauter

## Abstract

Network neuroscience models the brain as interacting elements. However, a large number of elements imply a vast number of interactions, making it difficult to assess which connections are relevant and which are spurious. Zalesky et al. (2010) proposed the Network-Based Statistics (NBS), which identifies clusters of connections and tests their likelihood via permutation tests. This framework shows a better trade-off of Type I and II errors compared to conventional multiple comparison corrections. NBS uses General Linear Hypothesis Testing (GLHT), which may underestimate the within-subject variance structure when dealing with longitudinal samples with a varying number of observations (unbalanced samples). We implemented NBR, an R-package that extends the NBS framework adding (non)linear mixed-effects (LME) models. LME models the within-subject variance in more detail, and deals with missing values more flexibly. To illustrate its advantages, we used a public dataset of 333 human participants (188/145 females/males; age range: 17.0-28.4 y.o.) with two (n=212) or three (n=121) sessions each. Sessions include a resting-state fMRI scan and psychometric data. State anxiety scores and connectivity matrices between brain lobes were extracted. We tested their relationship using GLHT and LME models for balanced and unbalanced datasets, respectively. Only the LME approach found a significant association between state anxiety and a subnetwork that includes the cingulum, frontal, parietal, occipital, and cerebellum. Given that missing data is very common in longitudinal studies, we expect that NBR will be very useful to explore unbalanced samples.

**Significant Statement:** Longitudinal studies are increasing in neuroscience, providing new insights into the brain under treatment, development, or aging. Nevertheless, missing data is highly frequent in those studies, and conventional designs may discard incomplete observations or underestimate the within-subject variance. We developed a publicly available software (R package: NBR) that implements mixed-effect models into every possible connection in a sample of networks, and it can find significant subsets of connections using non-parametric permutation tests. We demonstrate that using NBR on larger unbalanced samples has higher statistical power than when exploring the balanced subsamples. Although this method is applicable in general network analysis, we anticipate this method being potentially useful in systems neuroscience considering the increase of longitudinal samples in the field.

## 1. Introduction

Network science models complex phenomena from a system perspective, and its application covers a large variety of disciplines (Barabasi, 2016; Gosak et al., 2018). Particularly, network science has been applied to study the brain (Telesford et al., 2011; Basset & Sporns, 2017), where its structural and functional properties had been modeled from the interaction of its inner elements (defined at multiple scales), i.e., the brain connectome (Sporns, 2011; Craddock et al., 2013). This approach has been highly influential in non-invasive neuroimaging projects (Biswal et al., 2010; Van Essen et al., 2013), with great interest in its application to study the whole brain, longitudinally, across several time-scales and exploring multiple conditions (Di Martino et al., 2014). In this regard, longitudinal projects are currently increasing in order to address the insights of brain development, maturation, and degeneration (Ewing et al., 2018; Somerville et al., 2018; Becht & Mills, 2020). Accordingly, new tools are needed to account for both the *connectome* and its inherent change over time.

The brain network is modelled as a set of nodes (elements), defined based on anatomy, function, or another relevant distinction; and their connections (edges), usually based on (undirected) structural or functional associations (e.g., covariance of their BOLD signal). In the case of undirected associations, a network composed of V nodes may have up to V*(V-1)/2 connections, and V^2^ in the case of directed associations. Thus, testing individual connections for statistical relevance (for example, the association with a health trait, or differences between groups) may result in low statistical power, needing to correct for the large number of multiple comparisons. The Network-Based Statistics (NBS; Zalesky et al., 2010) aims to assess the whole set of interactions of a network with an efficient trade-off between the control of false positives and statistical power. This method (described below in more detail) focuses on the statistical feasibility of clusters of connections within a network, and it has been widely used to study the connectome in health and disease (Zhang et al., 2011; Pannek et al., 2013; Knyazev et al., 2015; Baggio et al., 2018; Gracia-Tabuenca et al., 2020a, 2020b; Noble & Scheinost, 2020). Like other commonly used toolboxes in neuroscience, the original NBS toolbox (RRID:SCR_002454) is limited to general linear hypothesis testing. Although general linear models (GLM) are highly flexible to test relationships between several variables of interest, they are limited when handling longitudinal data. Generally, GLM handles within-subject variability by the means of the individual intercepts, as covariates. However, this may underestimate the within-subject variability of the remaining variables of interest, particularly, when dealing with unbalanced samples, i.e., subjects with a different number of observations (Friston et al., 2005; Chen et al., 2013; Winter et al., 2013). Suitable methods to fit longitudinal data include mixed-effects models (Lindstrom & Bates, 1990), through which, the within- and between-subject variability structure is more widely described, by including individual variability in the dependent variables and intercepts, and their covariance as well (Barr et al., 2013). Also, these models handle missing data in a more flexible way (Mallinckrodt et al., 2001; Krueger et al., 2004; López-Gutierrez et al., 2019).

Here, we introduce the implementation of mixed-effects models into the Network Based Statistics approach, in order to properly address statistical tests in unbalanced longitudinal (repeated measures) samples. In particular, we introduce the Network-Based R-statistics (NBR; RRID:SCR_019114) package that implements NBS with mixed-effects models. We opted for an R package because R is a free software environment available for many operating systems and platforms, with widely used implementations of mixed-effects. As a demonstration, NBR was applied to an unbalanced sample of 333 participants, with two to three sessions, for a total of 787 observations. Each session included an MRI scan plus psychometric assessment. We tested if the anxiety state score is related to the functional connectome, for both the largest balanced sample and the complete unbalanced dataset.

## 2. Methods

### 2.1 NBS: familywise error rates for network components

The Network-Based Statistics (NBS; RRID: SCR_002454) firstly implemented by Zalesky et al. (2010), aims to identified clusters of edges (i.e. components) within a network and calculates the familywise error (FWE) probabilities of them.

First, an *a priori* threshold is applied for every single edge, then those above the threshold that share nodes in common are considered components. These components are “valued” based on their (binary or weighted) sum of edges. Components FWE probabilities are computed based on a permutation test of the maximum statistic. That is, for every permutation, data is shuffled and network components are calculated. Then, the highest sum of edges for all components is stored as the maximum statistic. This process is repeated a great number of times to create a pseudo-aleatory null distribution of components sum. And based on this, the components FWE probabilities can be computed following how likely it is to achieve their observed sum by chance.

In addition, this approach can control the false positive rate by a non-parametric test based on the original data itself, while maintaining a higher statistical power than other mass multiple testing methods, such as false discovery rate (FDR; Benjamini & Hochberg, 1995).

### 2.2 NBR: the NBS for mixed-effects models

The present work aims to extend the NBS to mixed-effect models implemented in R (RCT, 2020; RRID: SCR_001905). NBS allows the application of edgewise general linear hypothesis testing (GLHT), however when dealing with longitudinal data, the approximation is to include individual intercepts as covariates and permutation is restricted to within individual balanced datasets, i.e., each individual must have the same number of observations. In this regard, mixed-effects models extend the linear model adding random-effects that can be interpreted as additional error terms, which take into account the correlation of observation within the same individual. Furthermore, they are more flexible handling a variable number of observations and/or missing values (Pinheiro & Bates, 2006).

A general linear mixed-effects model can be expressed as (Jiang, 2007):

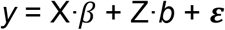

Where *y* stands for the response vector, X and Z are the fixed-effects and random-effects covariance matrices, while *β* and *b* are the fixed-effects and random-effects coefficients, respectively. *ε* stands for the error vector (for a detailed description of the random variance-covariance matrices and the parameters estimation see Pinheiro & Bates, 2006). In the context of a longitudinal study of brain connectomes and its relationship with a particular explanatory variable (e.g., phenotypic or genotypic), *y* will represent a single connectivity edge vector of *N* observations, grouped in *n* subjects. X will be a two-column design matrix including the intercept and the explanatory variable, while Z will be a 2x*n*-column matrix due to it includes the intercept and explanatory for each subject in separated columns, being zeros the positions that correspond to another subject.

NBR extends NBS by applying edgewise (non-)linear mixed-effects (LME) models through the R package ‘nlme’ (Pinheiro et al., 2017; RRID: SCR_015655). NBR functions allow the input of a 3D array of concatenated matrices along with the sample observations at the third dimension or a 2D matrix with every edge in the upper triangle of the matrices for each observation. The sample inference model is applied following the Wilkinson & Rogers (1973) notation, which is thoroughly used in R. If a group factor is specified in the random component of the LME, the randomization of the permuted data will be restricted to that factor even if it is unbalanced. The next steps follow those of the NBS: setting an *a priori* threshold, storing the observed components, generating a null distribution by permutation, and calculating the FWE p-value for each component (Figure 1). NBR (RRID: SCR_019114) can be downloaded via the CRAN (RRID: SCR_003005) in https://CRAN.R-project.org/package=NBR and it runs for Linux, Mac, and Windows platforms.

**Figure 1.**
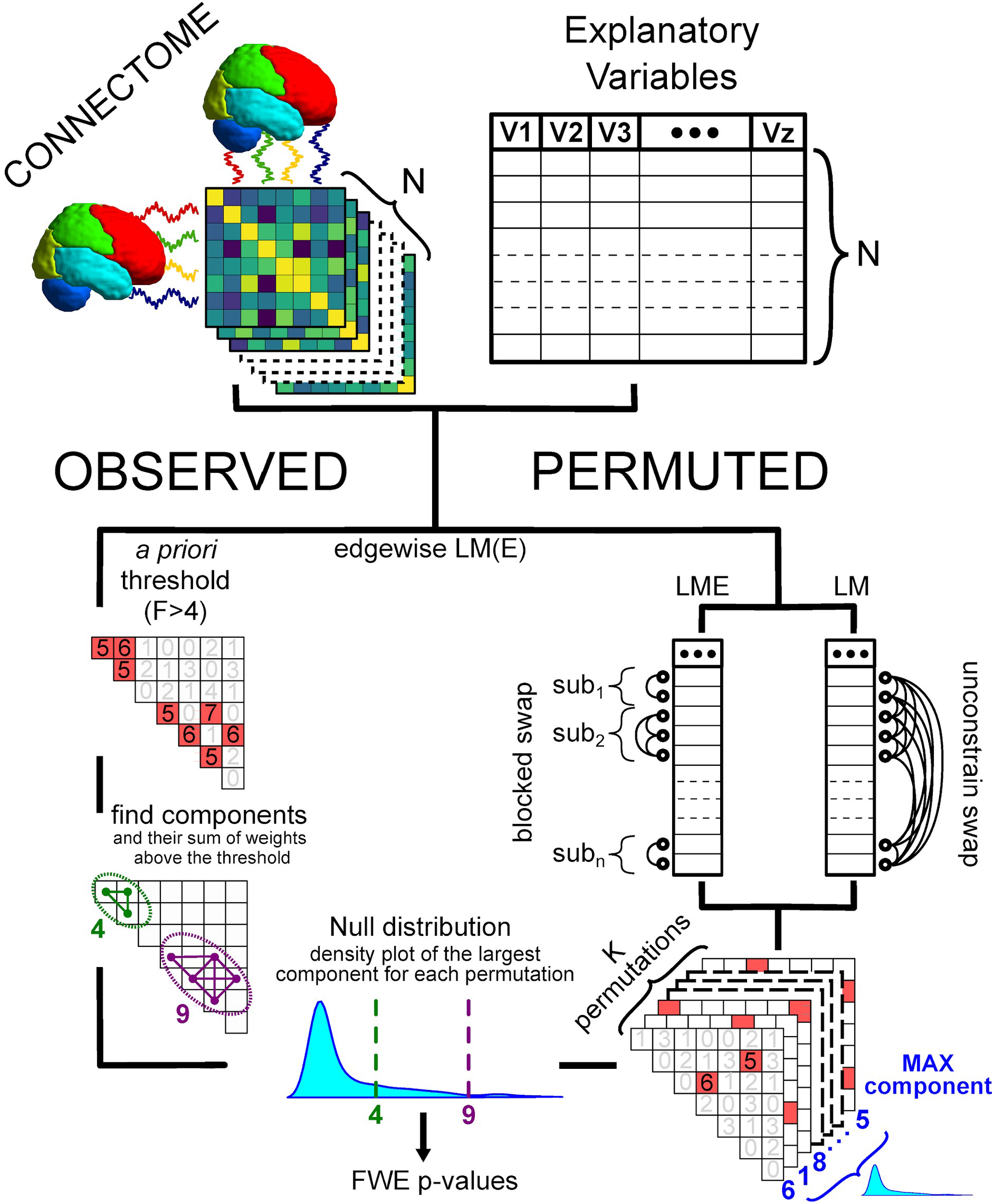
Network-based R-statistics diagram. Brain connectomes and variables of interest are the inputs. Then LM(E) is applied edgewise for the observed and permuted data. Finally, familywise error (FWE) p-values of the observed components are calculated comparing their edge sum respect to the maximum statistic null distribution obtained by K permutations.

### 2.3 Using NBR in a real-world longitudinal sample: the SLIM database

In order to illustrate a real case application, NBR was implemented in a public dataset. In particular, the Southwest University Longitudinal Imaging Multimodal (SLIM) database (Liu et al., 2017), which is publicly available for research through the International Data-sharing Initiative (INDI; http://fcon_1000.projects.nitrc.org/indi/retro/southwestuni_qiu_index.html). This is a large dataset (n = 595) focused on the neural basis of psychological traits, such as creativity and anxiety. The number of sessions varies between subjects between one, two, or three-time points. The average period between the first and second scans were 304.14 days, while between the second and the third sessions were 515 days. Each session included a multimodal MRI scan protocol and behavioral assessment using psychometric tests. Moreover, preprocessed functional connectivity matrices of resting-state fMRI scans are available. Scan parameters are available from Liu et al. (2017), and preprocessing steps followed the standard DPABI pipeline (Yan et al., 2016; RRID: SCR_010501).

Those subjects with at least two sessions were identified, resulting in a sample of 333 participants (145 males; age range: 17.0 - 28.4 y.o.), of which, 212 have two sessions and 121 have three sessions. Their corresponding functional connectivity matrices expressed as Fisher’s z-transformed correlation values, obtained with the Dosenbach160 brain segmentation (Dosenbach et al., 2010) were selected for further analyses. The connectivity values for each pair of regions between each pair of anatomical lobes (labeled by Watson, 2017) were further averaged, resulting in 8×8 connectivity matrices for each session.

We tested if psychometric anxiety scores are related to a specific brain network of interacting elements. In particular, the brain-behavior relationship taking into account longitudinal effects was evaluated taking the state component of the STAI psychometric test. The State-Trait Anxiety Inventory (STAI) is a Likert-like questionnaire that evaluates two types of anxiety: state or trait. Since the state component is less stable over time (Tian et al., 2016), it is a potential variable to test in relation with the functional connectome, including intra-individual effects in the models.

Several models were tested edgewise to illustrate the potential use of mixed-effects models in NBR compared to the common GLHT in NBS:

1. NBS-2tp: Given that the general linear model in NBS only allows balanced samples, the subsample of n = 211 subjects (98 male; 17.0 - 25.7 y.o. age range) with two sessions each (N = 422 observations) was explored with NBS.
2. NBS-3tp: Similar to the previous case but exploring the three available time points in the subsample of n = 53 subjects (25 male; 18.0 - 23.9 y.o. age range) with three sessions each (N = 159 observations).
3. NBR: LME of unbalanced two and three time points (n = 333 subjects). The complete subsample of subjects with at least two time points, resulting in a total of N = 787 observations.

Given that NBS tests only one-sided t-values, we opted to address the variability of the STAI by the means of F-test, setting F > 4 as an *a priori* threshold. This threshold was selected due to the probability of F = 4, given 1 and more than 100 degrees of freedom (which is the case for the three models) is close to 0.05.

### 2.4 Data and code availability

The summarized data and all the code implemented in this work is available in a public repository at https://github.com/BrainMapINB/NBR-SLIM. Present results were analyzed with R 3.4.4 and computed with an Intel Core i7-4790 CPU @ 3.60 GHz × 8 with Ubuntu 18.04.3 LTS 64-bit.

## 3. Results

The implementation of the network-based models in the SLIM dataset showed a component (cluster of connections) that involved frontal, parietal, and occipital lobes, cingulate, and cerebellum (Figure 2). However, after multiple comparison corrections only the component identified with NBR was statistically significant, i.e. under the nominal alpha (p_FWE_ < 0.05). Besides, not only the number of significant edges were higher for the NBR model, but its statistical strength was higher as well (Table 1). Lastly, the null distribution of the permuted original data showed lower variability for the NBR model (Figure 2).

**Table 1.** F-values for the edges above the *a priori* threshold (F>4) for each tested model.

|  | NBS-2tp | NBS-3tp | NBR |
| --- | --- | --- | --- |
| Cingulate-Cerebellum | 4.2238 | 7.5192 | 5.5079 |
| Frontal-Cerebellum |  | 4.0628 | 10.8919 |
| Frontal-Parietal |  | 6.2351 | 6.7341 |
| Occipital-Cigulate | 4.9174 |  | 12.0977 |

**Figure 2.**
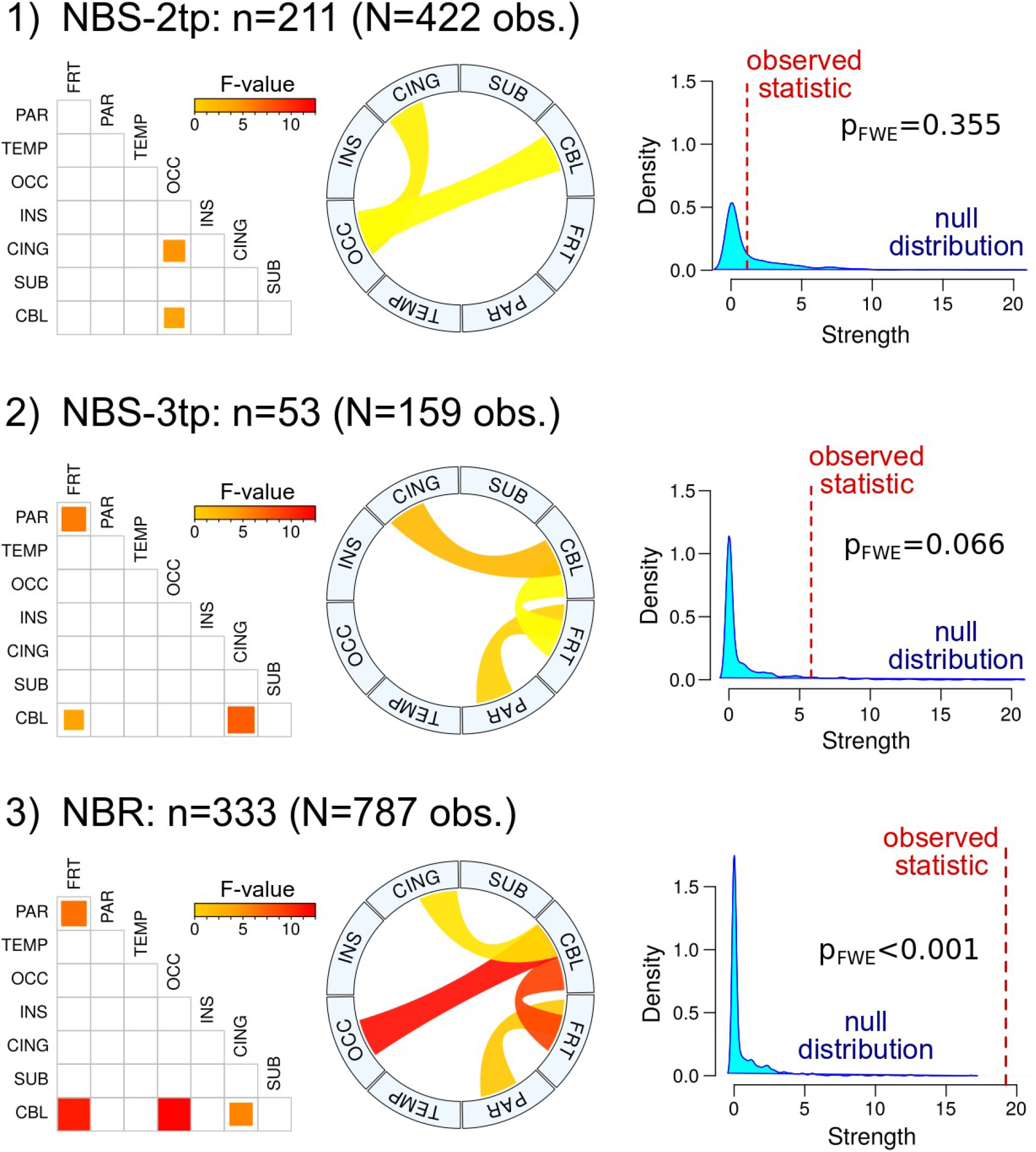
Implementation of network-based models: NBS-2tp (1), NBS-3tp (2), NBR longitudinal (3). For each model, its corresponding components are shown in pairwise (left) and circlesize (center) plots, next to the null distribution density plot of the largest component for 1000 permutations (right) with family-wise error p-value (p_FWE_). Red line in the density plots shows the weighted size of the component shown in the left. Abbreviations: frontal (FRT), parietal (PAR), temporal (TEMP), occipital (OCC), insula (INS), cingulate (CING), subcortical (SUB), and cerebellum (CBL).

## 4. Discussion

This work showed a new toolbox, NBR, designed to analyze longitudinal connectome samples, specifically, implementing mixed-effects models for Network-Based Statistics (NBS). Also, when applying it to a longitudinal sample, NBR was able to identify, beyond chance, a functional component of interconnected brain areas that are related to a varying psychological feature.

Regarding the NBR application example, this approach was able to identify a component of interacting brain regions that covariates with the anxiety state scores over time. Those regions include the frontal, parietal, cingulate, occipital, and cerebellum. The unbalanced mixed-effects model was able to combine the two clusters of connections that were identified when exploring the balanced datasets, i.e. the complete-cases designs. That is, NBR takes into account the covariance of the anxiety scores over time including both, two and three sessions. In addition, these results are congruent with previous studies taking a functional connectome approach that has found similar components in anxiety-related disorders (Maglanoc et al., 2019; Yang et al., 2019).

NBS relies on general linear hypothesis testing (GLHT), which may underestimate the within-subjects variability when dealing with longitudinal data. In this regard, NBR implements LME models to properly account for fixed and random variabilities. LME models can account for the within-subject variability of the intercepts, but also for the slopes, and their corresponding covariance. This may implicate a more complex model in terms of parameters, with its consequent decrease of statistical power, however, not considering this within-subject covariance structure may drive to an increase of false positive rates (Barr et al., 2013; Matuschek et al., 2017). Nevertheless, another relevant advantage of this approach is the possibility to explore all the observations in unbalanced samples, which in the application here explored resulted in a substantial increase in statistical power compared to using the possible balanced subsamples. Thus, the use of NBR not only allows for a more precise control of the within-subject variance, but allows the use of larger datasets when data is missing. Given the high costs of acquiring longitudinal datasets that imminently will suffer from missing data, NBR represents an ideal option to efficiently explore all the available observations using a network perspective. The main objective of NBR, as that of NBS, is to identify clusters of connections (components, or subnetworks) that satisfy a statistical test considering the available observations per subject.

We implemented NBR as an R package, because R is a free state-of-the-art statistical software available for Mac, Linux, and Windows operating systems. NBR can be run in R versions newer or equal to 2.10. Also, we opted to use the ‘nlme’ library to compute the LME instead of using other potential libraries that also implement these models because it is included in the default R base libraries. One potential limitation of the NBR is the high computation time. For instance, running the current example takes 55 minutes using the eight cores parallelization in the specified computer in the methods section. In contrast, running the NBS-2tp and NBS-3tp takes only 6 and 14 seconds, respectively. This occurs for two reasons, first the GLHT implies less computation steps than LME even with the same number of model parameters. Second, as the complexity of the within-subject covariance structure increases it does also the computing demand, for example, running the NBR-LME model but including just the within-subjects intercepts takes only 6 minutes to be computed. Besides, adding the anxiety scores along with the intercepts to the random effects (the ‘maximal model’) takes the aforementioned 55 minutes, because it estimates the random coefficients for the intercepts, slopes, and also their covariance. Although, the ‘maximal model’ is recommendable in terms of generalization (Barr et al., 2013), in particular cases it may have too many parameters, but the decision to select a more parsimonious model should be determined by the random effect structure that is supported by the data and not by the computation time (Matuschek et al., 2017). However, in order to lighten the computation time, NBR allows multi-core computing parallelization based on the 'parallel' library, which is also included in the R base libraries.

Lastly, NBR has additional features. Mixed-effects models may output different degrees of freedom for fixed and random variables of the model, for this regard, NBR allows setting the *a priori* threshold to define the clusters of connections, based on significance, i.e. the p-value, as an alternative to the use of the F, T or Z-values. The package also includes HTML vignettes with other examples and datasets, available in the CRAN repository (https://cran.r-project.org/web/packages/NBR/vignettes/).

## 5. Conclusion

Network-Based R-statistics is a new package to explore statistical inferences in longitudinal connectomes, including but not limited to brain-behavior relationships. Its suitability to fit within- and between-subject variability, even in unbalanced samples, makes NBR an appropriate method to address the upcoming growth of longitudinal datasets in neuroscience.

## Acknowledgements

We are extremely grateful to the International Neuroimaging Data-sharing Initiative, and particularly to the Southwest University Longitudinal Imaging Multimodal (SLIM) Brain Data Repository for such an important open data effort. We also thank Fernando A. Barrios for his valuable comments, Leopoldo Gonzalez-Santos for his technical support. Zeus Gracia Tabuenca is a doctoral student at the “Programa de Doctorado en Ciencias Biomédicas, Universidad Nacional Autónoma de México (UNAM)” and received a fellowship (330142) from “Consejo Nacional de Ciencia y Tecnología” (CONACYT). This research was partially supported by grant UNAM-DGAPA-PAPIIT IN212219 to SA. CONACYT and

DGAPA had no role in study design, data collection, analyses nor writing the manuscript. This work would not be possible without the guidance of the handbook *R packages* (https://r-pkgs.org/; Wickham, 2015) and the R community.

